# Experiment-anchored abundance–coverage frontiers define conditional RHSVV exposure requirements in TP53-missense tumors

**DOI:** 10.64898/2026.05.27.728141

**Authors:** Takehiro Ishikawa

## Abstract

Mutant p53 accumulation is established, but whether abundance can complement conditional RHSVV/PAb240-like exposure depends on the trade-off between target coverage and normal-tissue separation. We analyzed 809 tumors and 523 organ-matched adjacent-normal specimens from eight CPTAC cohorts. The 67.5% high-p53 fraction among quantified DNA-binding-domain missense tumors and the 2.59-fold paired tumor/normal median were treated only as dataset-specific secondary calibration. The design pool comprised 316 annotation-compatible missense tumors; 268 had quantified p53. Eligibility retained 179 tumors at cohort-normal P95, 159 at R≥2, 90 at R≥3, 54 at R≥4, 21 at R≥6, and 8 at R≥8, where R is abundance relative to the cohort-normal median. Normal stress used a prespecified 2.1-fold total-p53 abundance multiplier from a vinculin-normalized, multi-time-point ionizing-radiation study in early-passage wild-type Mdm2+/+ mouse embryonic fibroblasts; this source met all primary-anchor eligibility criteria. Under this anchor and a sample-level exposed-state mixture weight p=0.90, the full-data lowest passing target/background exposure grid point was q=2.00 pooled and q=2.50 worst cohort at P95, versus q=1.25 for both at R≥3; R≥3 covered 33.6% and is presented as a pragmatic exploratory intermediate, not an optimum. In 500 whole-pipeline resamples, all seven fixed cohorts were evaluable at R≥3 in 416 (83.2%); this is an evaluability fraction, not design or treatment success. Separate 0.05-margin-buffered fixed R≥3 bootstrap candidates used q=1.50 pooled and q=1.75 worst cohort and met the internal constraints in 678/1,000 pooled resamples and 628/863 evaluable worst-cohort resamples (628/1,000 overall; 137 were not cohort-evaluable). These post-selection internal sampling audits are neither therapeutic probabilities nor validation. The principal result is an experiment-anchored coverage–q frontier specifying conditional experimental requirements; native RHSVV exposure, tumor killing, and safety remain unmeasured.

## Introduction

DNA-damage signaling can stabilize wild-type p53 [1,2], while many TP53-missense tumors accumulate p53 constitutively. The mutant-accumulation phenotype is already supported by large proteomic resources and induced-proximity studies [3,4]; reproducing it in another complete-case cohort is calibration rather than the principal novelty. The translational question is whether an abundance-defined indication can leave sufficient separation from normal wild-type-p53 cells under experimentally observed stress responses to support a linked therapeutic activity.

PAb240 recognizes a conformation-sensitive epitope centered on RHSVV at residues 213-217 of the p53 DNA-binding domain [5-7]. Accessibility is not established by sequence retention alone and need not be universal among TP53 mutants. Engineered antibody fragments establish intracellular recognition of RHSVV or FRHSVV [8-10]. Abundance and accessibility therefore cannot be assumed to behave as two independently observed Boolean locks.

We used consistently processed CPTAC proteomic and mutation matrices from eight cohorts with adjacent-normal specimens [11-13]. Their role was not to estimate a universal tumor-to-normal fold change. Instead, the observed abundance distributions supplied a normal-anchored substrate for mapping how progressively stricter abundance eligibility changes target coverage, cohort representation, and the RHSVV exposure performance required under experimentally anchored normal p53-abundance stress.

The central question was: among sequence-compatible TP53-missense tumors, where is the coverage– RHSVV-requirement frontier when normal wild-type-p53 stress is calibrated to a qualified experimental source? The 2.1-fold Pant–Lozano WT-MEF ionizing-radiation peak was prespecified as the sole operational anchor under the source-eligibility criteria described below. Because CPTAC does not measure native RHSVV accessibility, every result is conditional on a sample-level exposure mixture and its assumed independence from abundance.

## Methods

### Study design and context of use

The study combined secondary normal-anchored description of p53 abundance with a prespecified abundance-eligibility sweep, one experimentally anchored normal-abundance stress scenario, and exact empirical monotone-cutpoint analysis. The anchor-selection criteria and all eligibility landmarks were disclosed. R≥3 was highlighted for focused reporting as a pragmatic intermediate point on the coverage– requirement curve, not as an optimized or validated biomarker threshold. The context of use was hypothesis prioritization and experimental specification, not prediction of clinical efficacy or toxicity.

### CPTAC cohorts and input matrices

Processed CPTAC pan-cancer matrices were obtained from LinkedOmicsKB Release 1, data freeze v1.2 [11,12]. The analysis included clear-cell renal cell carcinoma (CCRCC), colon adenocarcinoma (COAD), head and neck squamous cell carcinoma (HNSCC), lung squamous cell carcinoma (LSCC), lung adenocarcinoma (LUAD), ovarian carcinoma (OV), pancreatic ductal adenocarcinoma (PDAC), and uterine corpus endometrial carcinoma (UCEC), the eight cohorts with tumor and adjacent-normal proteomic matrices suitable for normal-anchored analysis.

Inputs comprised gene-level log2 reference-intensity-normalized tumor and normal proteomics, somatic mutation MAFs, binary mutation matrices, and case lists. The p53 feature was TP53/ENSG00000141510. Identifiers were harmonized, but raw cross-cohort intensities were not pooled; calibration and thresholding were performed within cohort, followed by stratified or cohort-adjusted analyses.

### TP53 alteration annotation and analytic groups

MAF records were restricted to protein-altering consequences. A tumor was classified as “protein-altering TP53 mutation detected” when supported by a qualifying MAF record or the binary mutation matrix. An evaluable zero binary call without a qualifying MAF record was classified as “no protein-altering TP53 mutation detected,” not as wild type, because germline status, copy-number state, loss of heterozygosity, and all structural lesions were not established.

Alterations were annotated for missense, DNA-binding-domain (DBD; residues 94–292) missense, truncating, splice, in-frame, direct RHSVV-region substitution, and truncation predicted to remove the RHSVV-containing sequence. Sample-level flags preserved co-occurring alterations.

### Missing p53 measurements

Missing p53 abundance was neither assigned zero nor imputed. Abundance comparisons used quantified complete cases. For every eligibility landmark, coverage was reported against the quantified molecular pool and with all-case lower and upper bounds that classified every unquantified molecularly eligible tumor as ineligible or eligible, respectively. Detection rates and the earlier high-p53 descriptive bounds were retained as secondary missingness audits.

### Within-cohort normal anchoring and high-p53 classification

Let y_i_c be the processed log2 p53 intensity for specimen i in cohort c, and let m_c be the median among quantified adjacent-normal specimens in that cohort:

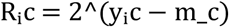

R entered the exact analysis without additional scaling or nonlinear transformation.

The cohort-normal 95th percentile was retained as a dataset-relative landmark. The indication sweep also used fixed R thresholds of 2, 3, 4, 6, and 8 so enrichment magnitude was explicit and comparable across cohorts. R≥3 was designated for focused reporting as a pragmatic intermediate between coverage and required q; no claim of optimality was attached to it. Eligibility threshold a and pharmacodynamic response cutpoint T were distinct: a selected the target population, whereas T was optimized only within a modeled exposure condition.

### Molecular eligibility, abundance eligibility, and coverage

The molecular pool comprised TP53-missense tumors whose RHSVV-containing sequence was compatible by the available mutation annotation. Variants directly affecting residues 213-217 and truncating co-alterations predicted to remove that sequence were excluded. This sequence filter is annotation-based, not a native-exposure assay. Each abundance landmark was applied to the same molecular pool. The seven cohorts with at least three P95-eligible tumors—COAD, HNSCC, LSCC, LUAD, OV, PDAC, and UCEC—defined the fixed reference set for cross-threshold worst-cohort comparison. A landmark or resample was non-evaluable for that analysis if any of the seven had fewer than three eligible tumors.

### Statistical analysis

High-p53 fractions were summarized with Wilson 95% confidence intervals. Continuous abundance was analyzed with cohort-specific Mann–Whitney tests and a cohort-adjusted linear model using HC3 standard errors. High-p53 classification was compared by mutation status with a cohort-stratified Cochran–Mantel–Haenszel analysis. Patient-matched log2 tumor-minus-normal differences were assessed with a 4,000-resample bootstrap confidence interval and two-sided Wilcoxon signed-rank test. Where multiple prespecified cohort or group comparisons formed a test family, P values were adjusted by the Benjamini–Hochberg procedure.

### Experimentally anchored normal p53-abundance stress

For the primary analysis, an eligible operational normal-stress anchor was required to provide (i) a wild-type-p53, non-neoplastic, normal-relevant system; (ii) direct measurement of total p53 protein abundance, the modeled quantity; (iii) ionizing-radiation exposure; (iv) a complete, traceable multi-time-point series with an explicit numerical peak; (v) loading-control normalization; and (vi) a directly reported fold change that could be used without selecting a range endpoint or rederiving an approximate value. We selected the Pant–Lozano early-passage Mdm2+/+ WT-MEF experiment because it met all six criteria. Its vinculin-normalized 0–10-h series after 6-Gy ionizing radiation peaked at 2.1-fold at 6 h [14]. We therefore used 2.1-fold as the sole operational normal-stress anchor and applied it to adjacent-normal p53 abundance. This source-level choice defines a traceable reporting baseline; it is not asserted to be a universal, most likely, or safety-conservative human normal-tissue response.

### Latent RHSVV exposure variables

CPTAC did not measure RHSVV accessibility. The model therefore swept the sample-level mixture weight p for eligible target observations in a high-exposure state and the target/background exposure contrast q=f_target/f_background. With f_target=1 and f_background=1/q, the common scale was absorbed by The implementation analytically averaged high- and background-exposure response curves and assumed the exposure state was independent of measured R. Thus, p is not identified as patient prevalence, within-tumor cellular fraction, or time spent in an exposed state.

Adjacent normals and tumors with no protein-altering TP53 mutation detected used the background state 1/q. For stressed normals, R was replaced by sR with s fixed at 2.1 from the selected operational anchor. No empirical joint R-by-exposure distribution was available. The model therefore assigns neither universal exposure to mutant tumors nor zero exposure to normal cells.

### Exact monotone-response analysis

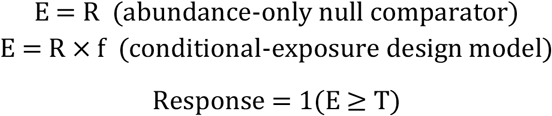

Under the selected 2.1-fold anchor, candidate T values were evaluated for each eligibility, mixture weight p, and contrast q combination. Candidate T values were the unique observed effective signals plus boundary sentinels. The selected T maximized the minimum margin to the prespecified constraints, with selection restricted to passing cutpoints whenever one existed. Reported contrasts are the lowest passing evaluated grid values, not continuous mathematical minima.

Targets were evaluated as a prevalence-weighted mixture of high- and background-exposure response curves. Pooled efficacy used all eligible targets. The stricter mode used the minimum response across the fixed set of seven P95-evaluable cohorts, each requiring at least three eligible samples at the analyzed landmark. This fixed-set rule prevents apparent improvement caused solely by loss of difficult cohorts.

### Parameter grids and passing criteria

Under the selected 2.1-fold anchor, p was evaluated from 0.50 through 1.00 in 0.05 increments across all six abundance landmarks. Both sampling audits used p=0.90. Contrast used quarter-step values from 1.00 through 20.00 and geometrically spaced values above 20 through 1,000. Passing required applicable target response at least 0.80, worst-cohort baseline adjacent-normal response at most 0.05, and worst-cohort abundance-stressed-normal response at most 0.10. q=1 denotes no exposure contrast and is equivalent to abundance-only thresholding.

### Sampling analyses and reproducibility checks

Sampling sensitivity was evaluated at p=0.90 under the primary 2.1-fold anchor. First, 500 within-cohort whole-pipeline resamples re-estimated normal anchors, abundance eligibility, q, and T for P95, R≥2, R≥3, and R≥4. Tumors were resampled within cohort; normals were resampled within cohort and quantified/missing strata. Fixed-set worst-cohort results were evaluable only when all seven reference cohorts retained at least three eligible tumors; the evaluable fraction measures representation, not design success. Second, the same fixed-candidate rule was applied to every landmark and efficacy mode: on the full data, select the smallest evaluated q admitting a T with at least 0.05 minimum margin, then hold p, q, and T fixed in 1,000 within-cohort resamples. We report response-constraint passes among all resamples and, for worst-cohort designs, among evaluable resamples, while separating cohort attrition. Both analyses are internal post-selection sampling audits, not held-out or external validation and not estimates of therapeutic success.

## Results

### Secondary dataset calibration reproduced the established missense-accumulation phenotype

The eight cohorts contained 809 tumors and 523 adjacent-normal specimens; p53 was quantified in 592 tumors and 371 adjacent-normal specimens, including 327 patient-matched pairs (Figure 1). Mutation status was evaluable in 804 tumors. These complete-case descriptions served to verify that the selected CPTAC release recapitulated known p53 accumulation and to define its normal-anchored R distributions.

**Figure 1.**
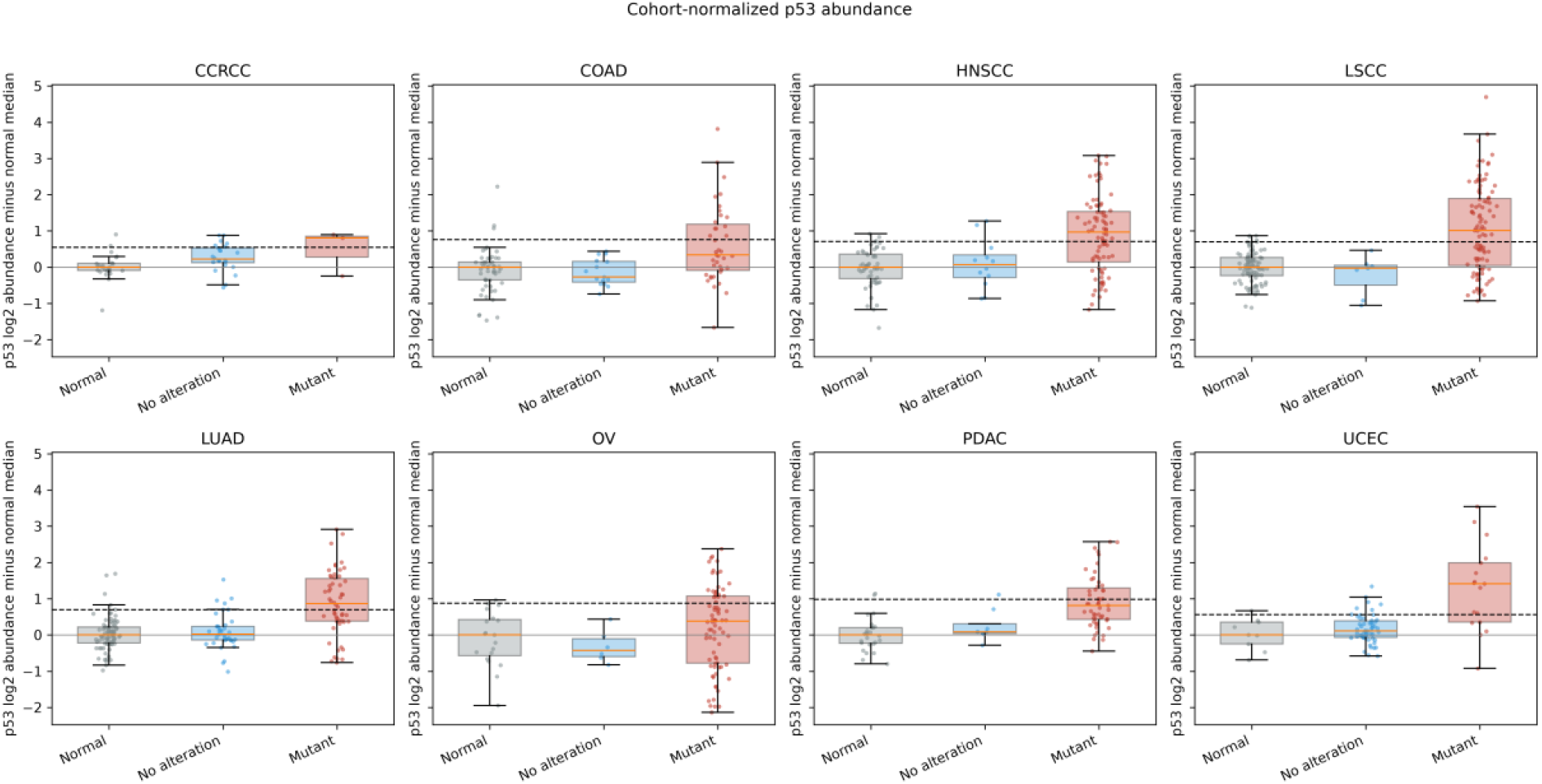
Cohort-normalized p53 abundance in adjacent-normal specimens and tumors with or without a detected protein-altering TP53 mutation. Values are log2 intensity minus the cohort-specific adjacent-normal median; dashed lines mark cohort-normal P95 thresholds. This is dataset-specific secondary calibration, not the primary design result.

Among quantified samples, 187 of 277 DBD-missense tumors (67.5%) exceeded the cohort-normal P95, compared with 23 of 163 tumors with no protein-altering TP53 mutation detected (14.1%) and 25 of 371 leave-one-out classified adjacent normals (6.7%; Table 1). These are complete-case proportions in this release, not population prevalences.

**Table 1.** Secondary complete-case calibration of high-p53 fractions under cohort-specific adjacent-normal P95 thresholds. Adjacent normals use leave-one-out thresholds; bounds assign every unquantified tumor to the low- or high-p53 category.

| Group | High/quantified n/N | High (%) | Wilson 95% CI (%) | All-sample bounds (%) |
| --- | --- | --- | --- | --- |
| Protein-altering TP53 mutation detected | 206/425 | 48.5 | 43.8–53.2 | 40.2–57.2 |
| TP53 missense | 191/285 | 67.0 | 61.4–72.2 | 57.2–71.9 |
| TP53 DBD missense | 187/277 | 67.5 | 61.8–72.8 | 57.9–72.1 |
| Other protein-altering | 15/140 | 10.7 | 6.6–16.9 | 8.4–29.8 |
| No protein-altering TP53 mutation detected | 23/163 | 14.1 | 9.6–20.3 | 7.9–52.1 |
| Adjacent normal, leave one out | 25/371 | 6.7 | 4.6–9.8 | — |

DBD, DNA-binding domain. Fractions are specific to quantified complete cases in this CPTAC release. Bounds are descriptive extremes, not probability intervals; the adjacent-normal row is a calibration check, not an independent validation set.

After cohort adjustment, protein-altering mutation status was associated with 0.831-log2 higher p53 abundance than no protein-altering mutation detected (approximately 1.78-fold; HC3 Wald z=9.77; P=1.49×10^-22). This comparison describes the dataset used for the design map rather than establishing a new accumulation mechanism.

In the paired complete cases, 159 DBD-missense tumors had a median 2.59-fold bulk tumor-to-adjacent-normal increase, whereas 83 tumors with no protein-altering TP53 mutation detected had a median 1.09-fold change and no significant paired increase (Table 2). The 2.59-fold value includes low-abundance tumors and is not the median contrast of the abundance-selected indication or a cell-level therapeutic window.

**Table 2.** Secondary dataset-specific patient-matched bulk tumor-to-adjacent-normal p53 abundance. Confidence intervals used 4,000 bootstrap resamples; P values are from two-sided Wilcoxon signed-rank tests.

| Analysis group | Pairs | Median log2 Δ | Median fold | Bootstrap 95% CI (log2) | Wilcoxon P | FDR |
| --- | --- | --- | --- | --- | --- | --- |
| All quantified pairs | 327 | 0.495 | 1.409 | 0.407 to 0.637 | $3.82 \times 10^{-25}$ | $3.72 \times 10^{-24}$ |
| Protein-altering mutation detected | 241 | 0.757 | 1.690 | 0.598 to 1.003 | $4.43 \times 10^{-27}$ | $1.73 \times 10^{-25}$ |
| TP53 missense | 164 | 1.371 | 2.587 | 1.059 to 1.644 | $1.42 \times 10^{-26}$ | $2.77 \times 10^{-25}$ |
| TP53 DBD missense | 159 | 1.373 | 2.590 | 1.017 to 1.649 | $8.29 \times 10^{-26}$ | $1.08 \times 10^{-24}$ |
| No protein-altering mutation detected | 83 | 0.125 | 1.091 | −0.096 to 0.234 | 0.522 | 0.566 |

FDR values use Benjamini-Hochberg correction across the full paired-analysis family. The fold estimates are bulk paired summaries, not abundance-selected therapeutic contrasts.

### Prespecified abundance enrichment exposed a coverage and cohort-representation trade-off

The annotation-defined molecular pool contained 316 sequence-compatible missense tumors, of which 268 had quantified p53. The P95 landmark retained 179 tumors, corresponding to 66.8% complete-case coverage; the all-case lower and upper bounds were 56.6% and 71.8%. P95 retained at least three tumors in seven cohorts; CCRCC retained two and was not in the fixed comparison set (Figure 2).

**Figure 2.**
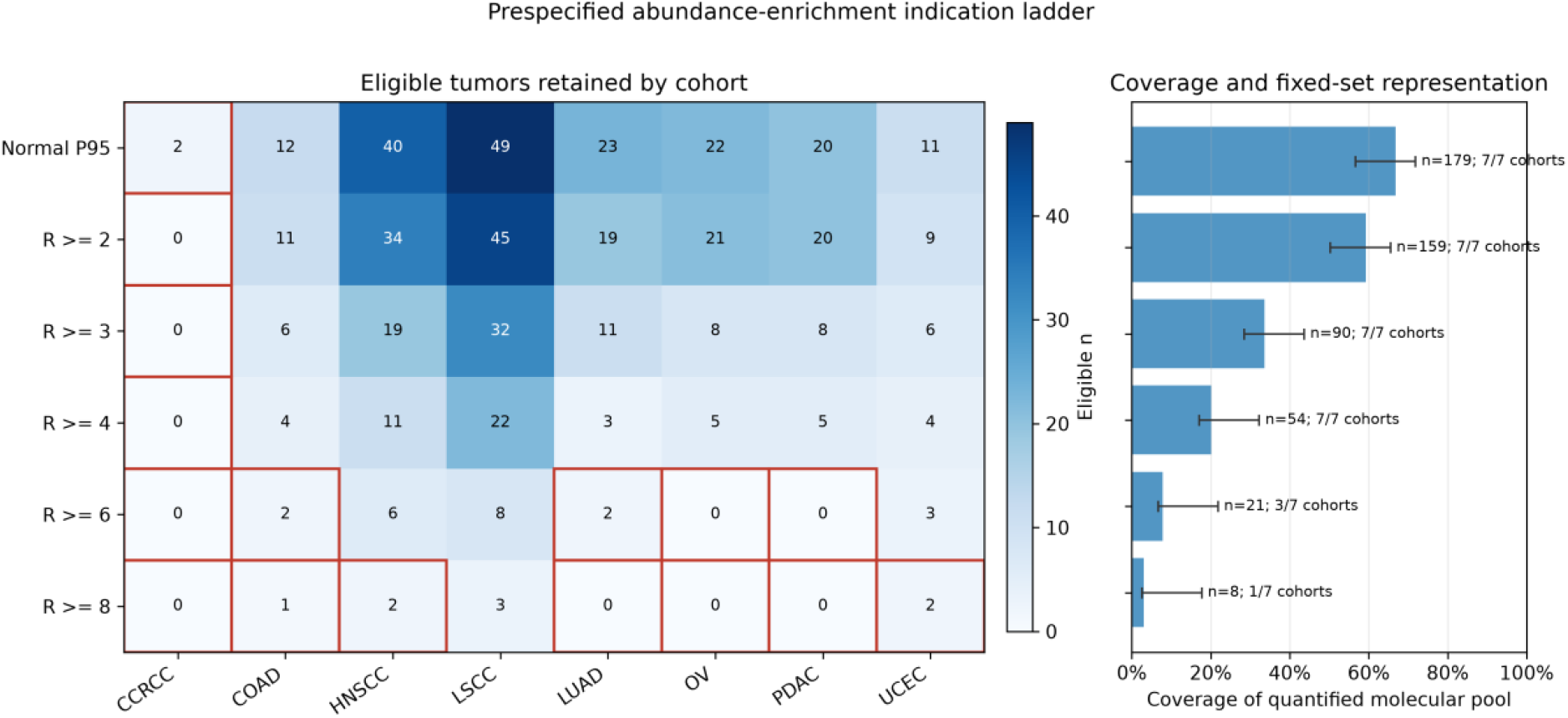
Prespecified abundance-enrichment indication ladder. The heat map gives eligible tumors by cohort; red outlines mark cells with fewer than three tumors. Bars show coverage of 268 quantified molecularly eligible tumors, labels show fixed reference cohorts retained, and whiskers span all-case lower and upper bounds obtained by assigning all 48 unquantified molecularly eligible tumors as ineligible or eligible. R≥3 is highlighted as a pragmatic exploratory intermediate, not an optimized or validated cutoff.

Fixed enrichment thresholds retained 159 tumors at R>=2 (59.3% complete-case coverage), 90 at R>=3 (33.6%), 54 at R>=4 (20.1%), 21 at R>=6 (7.8%), and 8 at R>=8 (3.0%). P95 through R>=4 retained all seven fixed-set cohorts. R>=6 retained three, and R>=8 retained one; those two landmarks were therefore not interpreted under the fixed-set worst-cohort criterion.

### Experiment-anchored abundance enrichment defined a coverage–contrast frontier

Under the primary 2.1-fold normal-abundance stress anchor, abundance alone (q=1) did not pass at P95, R≥2, or R≥3. It first passed both pooled and fixed-set worst-cohort criteria at R≥4, which covered 54 of 268 quantified molecularly eligible tumors (20.1%). The more stringent R≥6 and R≥8 landmarks covered only 7.8% and 3.0% and no longer retained the seven fixed cohorts.

Across the full 11-point p grid under the selected 2.1-fold anchor, the lowest evaluated p admitting any grid-passing q was 0.75 pooled and 0.80 fixed-set worst cohort at P95, 0.70 and 0.75 at R≥2, and 0.50 and 0.70 at R≥3. R≥4 passed at q=1 across all evaluated p values, so p did not affect that abundance-only case. Complete statuses and q and T values are reported in Supplementary Table S1. At p=0.90, the lowest evaluated passing q was 2.00 pooled and 2.50 fixed-set worst cohort at P95; 1.75 and 2.00 at R≥2; 1.25 and 1.25 at R≥3; and 1.00 and 1.00 at R≥4 (Table 3; Figure 3). R≥3 retained 90 tumors (33.6%) and all seven cohorts in the full dataset, making it a pragmatic exploratory intermediate, not an optimum or validated cutoff.

**Table 3.**
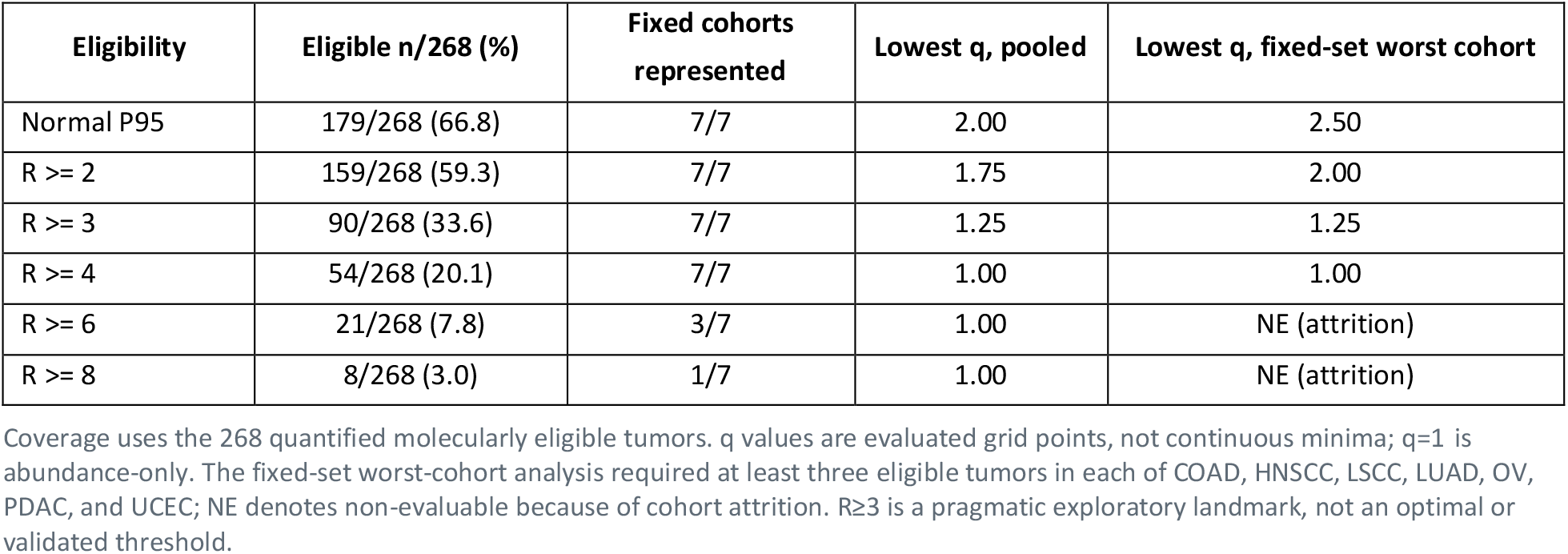
Abundance eligibility, coverage, cohort representation, and lowest evaluated passing RHSVV target/background exposure contrast at p=0.90 under the selected Pant–Lozano WT-MEF ionizing-radiation anchor (2.1-fold normal p53-abundance stress). This is the p=0.90 reporting slice used for sampling audits; all 11 evaluated p values are reported in Supplementary Table S1. Passing required target response at least 0.80, worst-cohort baseline-normal response at most 0.05, and worst-cohort abundance-stressed-normal response at most 0.10.

| Eligibility | Eligible n/268 (%) | Fixed cohorts represented | Lowest q, pooled | Lowest q, fixed-set worst cohort |
| --- | --- | --- | --- | --- |
| Normal P95 | 179/268 (66.8) | 7/7 | 2.00 | 2.50 |
| $R \geq 2$ | 159/268 (59.3) | 7/7 | 1.75 | 2.00 |
| $R \geq 3$ | 90/268 (33.6) | 7/7 | 1.25 | 1.25 |
| $R \geq 4$ | 54/268 (20.1) | 7/7 | 1.00 | 1.00 |
| $R \geq 6$ | 21/268 (7.8) | 3/7 | 1.00 | NE (attrition) |
| $R \geq 8$ | 8/268 (3.0) | 1/7 | 1.00 | NE (attrition) |

Coverage uses the 268 quantified molecularly eligible tumors. $q$ values are evaluated grid points, not continuous minima; $q=1$ is abundance-only. The fixed-set worst-cohort analysis required at least three eligible tumors in each of COAD, HNSCC, LSCC, LUAD, OV, PDAC, and UCEC; NE denotes non-evaluable because of cohort attrition. $R \geq 3$ is a pragmatic exploratory landmark, not an optimal or validated threshold.

**Figure 3.**
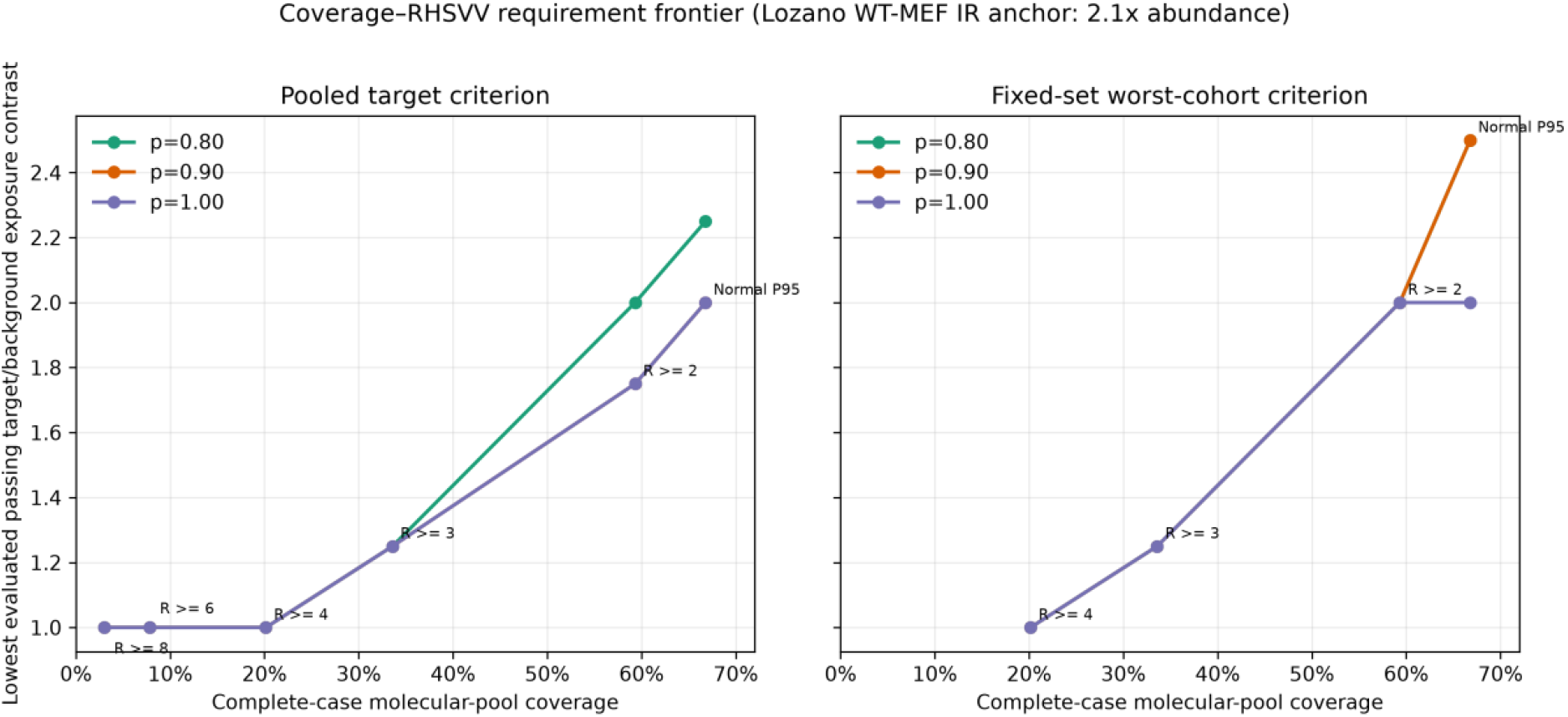
Coverage–RHSVV requirement frontier under the primary 2.1-fold Lozano WT-MEF ionizing-radiation abundance anchor. Points show the lowest evaluated passing q at sample-level exposed-target mixture weights p=0.80, 0.90, and 1.00. These three readability-selected slices come from the full 11-point p grid; complete results are provided in Supplementary Table S1. The right panel is restricted to landmarks retaining all seven fixed reference cohorts. Lines connect discrete landmarks and are not continuous biological functions; R≥3 is an exploratory intermediate, not an optimum.

**Figure 4.**
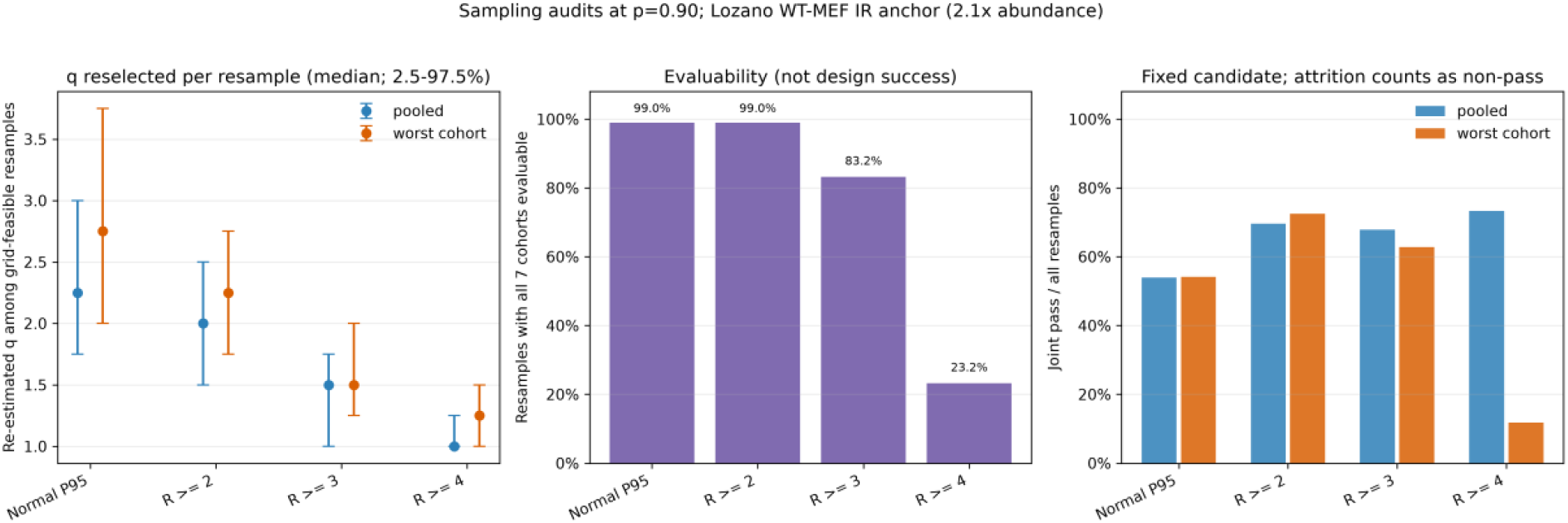
Internal sampling audits at p=0.90 under the primary 2.1-fold Lozano WT-MEF ionizing-radiation abundance anchor. Left: median and 2.5th-97.5th percentiles of q reselected in each of 500 whole-pipeline resamples, among grid-feasible resamples; worst-cohort summaries are additionally conditional on fixed-set evaluability. Middle: the fraction of those 500 resamples retaining at least three eligible tumors in every fixed reference cohort; 416/500 (83.2%) at R≥3 is evaluability only, not design success. Right: the fraction of all 1,000 resamples meeting the response constraints with the full-data-selected p, q, and T held fixed; worst-cohort bars count cohort-attrition resamples as non-passes, and conditional denominators are reported in the text. These post-selection internal audits are neither therapeutic success probabilities nor validation.

### Sampling audits separated q sensitivity from cohort evaluability

In 500 whole-pipeline resamples at p=0.90 under the 2.1-fold anchor, median re-estimated q (2.5th-97.5th percentiles) at P95 was 2.25 (1.75-3.00) pooled and 2.75 (2.00-3.75) worst cohort. At R≥3, the corresponding values were 1.50 (1.00-1.75) and 1.50 (1.25-2.00). Pooled q was grid-feasible in 500/500 R≥3 resamples, irrespective of fixed-cohort retention. All seven fixed cohorts were evaluable in 416/500 R≥3 resamples (83.2%), and a worst-cohort grid solution existed in all 416 of those evaluable resamples. The 83.2% is solely a fixed-seven-cohort evaluability fraction; it is not a design-pass or treatment-success rate.

For the focused R≥3 fixed-design audit, the full-data pooled candidate used p=0.90, q=1.50, and T=3.132540; the worst-cohort candidate used p=0.90, q=1.75, and T=2.718309. Both had a full-data minimum margin of 0.05. With these values held fixed, pooled response constraints were met in 678/1,000 resamples (67.8%; Wilson 95% interval, 64.8%-70.6%). For the worst-cohort candidate, 863/1,000 resamples retained all seven cohorts; 628/863 evaluable resamples met the response constraints (72.8%; 69.7%-75.6%), equivalent to 628/1,000 when the 137 cohort-attrition resamples were counted separately as non-passes. These fractions describe post-selection stability under resampling; they are not probabilities of tumor killing, toxicity avoidance, or therapeutic success, and they do not constitute validation.

## Discussion

This analysis changes the emphasis from re-estimating a known phenotype to defining an experiment-anchored conditional indication design. The 67.5% P95 proportion and 2.59-fold paired median confirm that this CPTAC complete-case subset contains the established missense-accumulation signal [3,4], but both are dataset-specific secondary calibration. Bulk composition, tumor purity, platform normalization, normal comparator, and cohort mix affect their magnitude.

Abundance and RHSVV exposure are not two independent observed keys. The modeled quantity is the composite E=R×f: a normal cell with high R and background exposure can overlap a tumor with lower R and higher exposure. The analysis assumed the sample-level exposure mixture p was independent of abundance because their joint distribution was unavailable.

The central design insight is the coverage–q frontier. Under the selected 2.1-fold anchor, moving from P95 to R≥3 reduced the observed-sample q requirement from 2.00/2.50 to 1.25/1.25 for pooled/worst-cohort criteria, while coverage fell from 66.8% to 33.6%. We highlight R≥3 because it occupies a pragmatic intermediate position and retained all seven fixed cohorts in the full data—not because it was optimized or validated.

The decisive next experiment is a joint measurement, not another threshold search. Total p53 and native RHSVV accessibility should be measured in the same tumor and matched-normal specimens, ideally at single-cell resolution, so their empirical joint R-by-f distributions can be placed on the frontier. Total-p53 abundance should be measured after stress in wild-type-p53 normal models solely to calibrate the stress multiplier. A real binder must subsequently demonstrate intracellular target engagement and calibrated cell death before the dimensionless threshold model can inform efficacy or safety.

## Limitations

CPTAC measures total p53 abundance, not native RHSVV accessibility. p and q are swept assumptions, not observed biomarkers. The sample-level response mixture assumes exposure independent of R and cannot distinguish patient-level prevalence from within-tumor cellular heterogeneity or temporal exposure. The full p grid was evaluated under the selected 2.1-fold anchor; both sampling audits used p=0.90. Sequence compatibility does not establish native PAb240/RHSVV accessibility [5-7].

Adjacent-normal tissue is an organ-matched bulk reference, not healthy whole-body tissue. Field effects, tumor admixture, cell-type composition, and the eight-cohort organ scope limit safety interpretation.

p53 was unquantified in 48 of 316 molecularly eligible missense tumors and in 152 of 523 adjacent-normal specimens. Complete-case coverage may therefore be detection-biased despite the reported bounds. Sequence compatibility was inferred from available annotations; complex in-frame or splice effects on residues 213-217 may be incompletely captured. ‘No protein-altering mutation detected’ also does not establish germline, copy-number, structural, loss-of-heterozygosity, or purity status.

Threshold crossing is not observed cell killing, and the normal constraint is not clinical toxicity. All landmarks were analyzed on the same dataset. Disclosing the entire ladder limits hidden threshold selection, but the 500-resample whole-pipeline analysis and the 1,000-resample fixed-candidate analysis remain internal sampling audits rather than external or held-out validation. Cohort attrition is a representation problem and was separated from response-constraint failures. The cohort minimum of three is small, leaving wide uncertainty in cohort-specific tails.

The 2.1-fold anchor was selected for source fitness, not because it is universal or maximally conservative; it remains a mouse-MEF value from a specific 6-Gy ionizing-radiation experiment [14]. Direct quantitative measurements across relevant human normal cell types and clinically relevant insults are needed. The 5% and 10% worst-cohort normal limits are also highly discrete in small cohorts.

## Conclusion

The principal contribution is an experiment-anchored coverage–q frontier for an abundance-plus-RHSVV strategy. Under the selected 2.1-fold normal-abundance anchor at p=0.90, P95 covered 66.8% of the quantified molecular pool and required q=2.00 pooled and 2.50 fixed-set worst cohort; the pragmatic R≥3 intermediate covered 33.6% and required q=1.25 for both. R≥3 is neither optimal nor validated. These results specify falsifiable experimental requirements, not evidence of native RHSVV exposure, tumor-selective killing, clinical safety, or treatment success.

## Ethics

No new human participant, human tissue, or animal data were generated in this study.

## Data accessibility

Processed CPTAC pan-cancer data-freeze v1.2 matrices were obtained from LinkedOmicsKB Release 1 [11,12] at https://kb.linkedomics.org/download. The upstream CPTAC resource is available through the NCI Proteomic Data Commons [15]. The executable notebook, primary-anchor source documentation, derived frontier, and sampling-audit outputs for this revised manuscript are available on Zenodo at https://doi.org/10.5281/zenodo.21964384.

## Declaration of AI use

AI-assisted tools were used for language editing and programming assistance. All code, analyses, outputs, interpretations, claims, and final manuscript content were reviewed and approved by the human author.

## Author contributions

T.I: conceptualization, data curation, formal analysis, investigation, methodology, software, validation, visualization, writing—original draft, and writing—review and editing.

## Acknowledgements

Data used in this publication were generated by the Clinical Proteomic Tumor Analysis Consortium (NCI/NIH).

